# Y-chromosome haplogroups from Hun, Avar and conquering Hungarian period nomadic people of the Carpathian Basin

**DOI:** 10.1101/597997

**Authors:** Endre Neparáczki, Zoltán Maróti, Tibor Kalmár, Kitti Maár, István Nagy, Dóra Latinovics, Ágnes Kustár, György Pálfi, Erika Molnár, Antónia Marcsik, Csilla Balogh, Gábor Lőrinczy, Szilárd Sándor Gál, Péter Tomka, Bernadett Kovacsóczy, László Kovács, István Raskó, Tibor Török

## Abstract

Hun, Avar and conquering Hungarian nomadic groups arrived into the Carpathian Basin from the Eurasian Steppes and significantly influenced its political and ethnical landscape. In order to shed light on the genetic affinity of above groups we have determined Y chromosomal haplogroups and autosomal loci, from 49 individuals, supposed to represent military leaders. Haplogroups from the Hun-age are consistent with Xiongnu ancestry of European Huns. Most of the Avar-age individuals carry east Eurasian Y haplogroups typical for modern north-eastern Siberian and Buryat populations and their autosomal loci indicate mostly unmixed Asian characteristics. In contrast the conquering Hungarians seem to be a recently assembled population incorporating pure European, Asian and admixed components. Their heterogeneous paternal and maternal lineages indicate similar phylogeographic origin of males and females, derived from Central-Inner Asian and European Pontic Steppe sources. Composition of conquering Hungarian paternal lineages is very similar to that of Baskhirs, supporting historical sources that report identity of the two groups.

## Introduction

Population history of the Carpathian Basin was profoundly determined by the invasion of various nomadic groups from the Eurasian Steppes during the Middle Ages. Between 400-453 AD the Huns held possession of the region and brought about a major population reshuffling all over Europe. Recent genetic data connect European Huns to Inner Asian Xiongnus ^1^, but genetic data from Huns of the Carpathian Basin have not been available yet, since Huns left just sporadic lonely graves in the region, as they stayed for short period. We report three Y haplogroups (Hg) from Hun age remains, which possibly belonged to Huns based on their archaeological and anthropological evaluation.

From 568 AD the Avars established an empire in the region lasting nearly for 250 years, until they were c, then their steppe-empire ended around 822 AD. In its early stage the Avar Khaganate controlled a large territory expanding from the Carpathian Basin to the Pontic-Caspian Steppes and dominated numerous folks including Onogur-Bulgars, which fought their independence in the middle 7th century and established the independent Magna Bulgaria state. Then the Avar Khaganate was shrunken, its range well corresponding to that of the succeeding Hungarian Kingdom. cand the Avar period left a vast archeological legacy with more than 80 thousand excavated graves in present-day Hungary. The Avar age remains are anthropologically extremely heterogeneous, with considerable proportion of Mongoloid and Europo-Monoloid elements reaching 20-30% on the Great Hungarian Plain ^2^, attesting that the Carpathian Basin witnessed the largest invasion of people from Asia during this period. Most individuals buried with rich grave goods show Mongoloid characters indicating inner Asian origin of the Avar elite, which is also supported by their artifact types, titles (e.g. khagan) and institutions recognized to be derived from Inner Asian Rourans. From the Avar period only a few mitochondrial DNA (mtDNA) data are available from two micro-regions, ^3,4^ which showed 15.3% and 6.52% frequency of East Eurasian elements. A recent manuscript described 23 mitogenomes from the 7^th^-8^th^ century Avar elite group ^5^ and found that 64% of the lineages belong to East Asian haplogroups (C, D, F, M, R, Y and Z) with affinities to ancient and modern Inner Asian populations corroborating their Rouran origin. Though the Avar Khaganate ceased to exist around 822 AD, but its population survived and were incorporated into the succeeding Hungarian state ^6^. It is relevant to note that none of the Hungarian medieval sources know about Avars ^7^, probably because they were not distinguished from the Huns as many foreign medieval sources also identified Avars with Huns, for example the Avars who were Christianized and became tax-payer vassals of the Eastern Frankish Empire were called as Huns in 871 ^8^.

Presence of the Hungarians in the Carpathian Basin was documented from 862 AD and between 895-905 they took full command of the region. The Hungarians formed a tribal union but arrived in the frame of a strong centralized steppe-empire under the leadership of prince Álmos and his son Árpád, who were known to be direct descendants of the great Hun leader Attila, and became founders of the Hungarian ruling dynasty and the Hungarian state. The Hungarian Great Principality existed in Central Europe from ca. 862 until 1000, then it was re-organized as a Christian Kingdom by King István I the Saint who was the 5^th^ descendant of Álmos ^9^.

Although neither the Avar, nor the Hungarian steppe-state was homogenous, their components were originated in the same cultural background of the Eurasian steppe-belt. During the 6^th^–7^th^ centuries Kutrighur, Sabir and later Onogur immigrating groups joined the Avars ^6^ which were all Hun-Turkic people and the Avars probably also had Hun ethnic components. The conquering Hungarians with their Turkic-speaking Kabar vassals integrated people of the former Avar state and among these people were the descendants of the Onogurs which were regarded proto-Hungarians by László ^10^. The Avar Khaganate also integrated Slavic and German groups, thus by the end of the 9^th^ century the majority of the population in the Carpathian Basin originated from the Hun-Turkic community of the Eurasian Steppe, complemented with Slavonic and German-speaking groups.

Our recent analysis of conquering Hungarian (hence shortened as Conqueror) mitogenomes revealed that the origin of their maternal lineages can be traced back to distant parts of the Eurasian steppe ^11^. One third of the maternal lineages were derived from Central-Inner Asia and their most probable ultimate sources were the Asian Scythians and Asian Huns, while the majority of the lineages most likely originated from the Bronze Age Potapovka-Poltavka-Srubnaya cultures of the Pontic-Caspian steppe. Population genetic analysis indicated that Conquerors had closest connection to the Onogur-Bulgar ancestors of Volga Tatars.

Although mtDNA data can be informative, since as far as we know none of the above nomadic invasions were pure military expeditions in which raiding males took local women, but entire societies with both men and women migrated together ^3^, however nomadic societies were patrilinealy organized, thus Y-chromosome data are expected to provide more relevant information about their structure and origin than mtDNA. So far 6 Y-chromosome Hg-s have been published from the Conquerors; ^12^ which revealed the presence of N1a1-M46 (previously called Tat or N1c), in two out of 4 men, while ^13^ detected two R1b-U106 and two I2a-M170 Hg-s.

In order to generate sufficient data for statistical evaluation and compare paternal and maternal lineages of the same population, we have determined Y-Hg-s from the same cemeteries whose mtDNA Hg-s were described in ^11^. As most Hungarian medieval chronicles describe the conquer as the “second incoming of the Hungarians” ^7^, genetic relations may exist between consecutive nomadic groups, first of all between elite groups. As the studied Conqueror cemeteries represent mainly the Conqueror elite, we supplemented our Y-Hg studies with samples from the preceding Avar military leader groups and available samples from the Hun period to test possible genetic relations. We also tested autosomal single nucleotide polymorphisms (SNP-s) associated with eye/hair/skin color and lactase persistence phenotypes as well as biogeographic ancestry.

## Results

We selected 168 phylogenetically informative Y chromosome SNP-s ^14^ defining all major Hg-s and the most frequent Eurasian sub-Hg-s, as well as the following 61autosomal SNP-s: 25 HirisPlex markers suitable for eye and hair color prediction ^15^, two SNP-s linked to adult lactase persistence ^16^ and 34 ancestry informative markers (AIMs) ^17^, as listed in Supplementary Table S1 and S2. DNA segments containing above SNP-s were enriched from the next generation sequencing (NGS) library of each sample with hybridization capture. Anthropological males were selected from the cemeteries, but as control we also included four females for which the presence of Y-chromosomal reads can reveal contamination. Analysis of women control samples revealed low contamination, having negligible Y-chromosomal reads but comparable autosomal reads to males (Supplementary Table S1 and S2) with one exception, KEF1/10936 turned out genetically a male despite its anthropological description as female. Negligible contamination levels for 26 of the 49 libraries had been demonstrated before ^11^ and low contamination is also inferred from unambiguous Hg classifications with very few contradicting SNP-s, most of which can be explained by postmortem ancient DNA modifications (Supplementary Table S1). Besides, MapDamage indicated typical ancient DNA transition patterns and library fragment size distributions also corresponded to aDNA (Supplementary Table S3). Moreover we detected a significant number of east Eurasian Hg-s which had a negligible chance to have been derived from a contaminating source of European researchers. We obtained low to medium average coverage of the targeted loci, (Supplementary Table S1, S2 and S3) and Y-Hg determination was consistent from 46 males (Supplementary Table S1) which is summarized in Table 1.

**Fig. 1.**
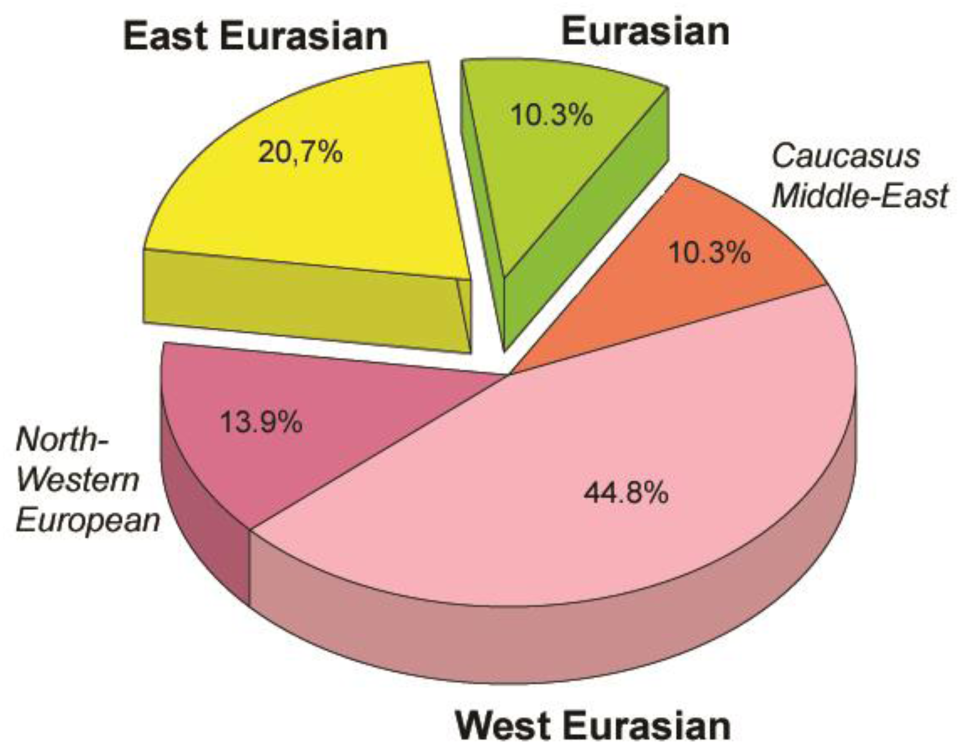
Distribution of the 29 Conqueror paternal lineages according to their phylogeographic origin. Data are summarized from Table 1.

**Table 1.** Y-Hg-s determined from 46 males grouped according to sample age, cemetery and Hg. Hg designations are given according to ISOGG Tree 2019. Grey shading designate distinguished individuals with rich grave goods, color shadings denote geographic origin of Hg-s according to Fig. 1. For samples K3/1 and K3/3 the innermost Hg defining marker U106* was not covered, but had been determined previously ^13^.

| LabID | Sample site/grave or accession number | Age | Innermost Hg defining marker covered | Possible innermost marker not covered | Y- Haplogroup |
| --- | --- | --- | --- | --- | --- |
| **Hun period** |  |  |  |  |  |
| Hun/1 | Singeorgiu de Mures/1 | V. century | M25 | - | Q1a2 |
| Hun/2 | Kecskemét-Mindszenti-dűlő/2785 | V. century | U106 | - | R1b1a1b1a1a1 |
| Hun/3 | Árpás-Szérűskert/1 | V. century | Z2124 | - | R1a1a1b2a2 |
| **early Avar period** |  |  |  |  |  |
| FGD/4 | Fajsz-Garadomb/4 | 630–650/660 | P44/M217 | - | C2 |
| PSZ/1 | Petőfiszállás/1 | 630–650/660 | P15 (xL1259) | L293 | G2a |
| SzO/540 | Szegvár-Oromdűlő/540 | 600–650/660 | M253 (xL22) | - | I1 |
| SzO/81 | Szegvár-Oromdűlő/81 | 600–650/660 | B197 | - | N1a1a1a1a3 |
| KFP/6 | Kunpeszér-Felsőpeszér út, Homokbánya/6 | 630–650/660 | Z35291 | - | N1a1a1a1a3 |
| KFP/31 | Kunpeszér-Felsőpeszér út, Homokbánya/30B | 630–650/660 | Z35291 | - | N1a1a1a1a3 |
| SzK/51 | Székkutas-Kápolnadűlő/51 | 630/650–660 | B197 | - | N1a1a1a1a3 |
| KB/300 | Kunbábony/300 (**khagan**) | 630–650/660 | M178 (xZ35159) | VL29, Z1936, L1034 | N1a1a |
| MM/58 | Makó-Mikócsa halom/56-58 | 568–630 | M178 (xVL29, xZ1936, xZ35159, xY9286, xB197, xM2004) | - | N1a1a |
| MM/227 | Makó-Mikócsa halom/218-227 | 568–630 | Z94 | Z2124 | R1a1a1b2a |
| DK/701 | Dunavecse-Kovacsos dűlő/701 | 630–650/660 | Z94 | Z2124 | R1a1a1b2a |
| **middle and late Avar period** |  |  |  |  |  |
| PV/72 | Pitvaros-Víztározó/72 | 650/660–700/710 | P44, M217 | - | C2 |
| KV/3369 | Kiskőrös-Vágóhídi dűlő/3369 | 650/660–700 | Z35291 | - | N1a1a1a1a3 |
| SzK/239 | Székkutas-Kápolnadűlő/239 | 650/660–700/710 | V13 | - | E1b1b1a1b1a |
| **Conqueror period** |  |  |  |  |  |
| K1/13 | Karos I/13 | 895- mid Xth c. | M215 | V13 | E1b1b |
| K1/1438 | Karos I/1438 | 895- mid Xth c. | M267 | - | J1 |
| K1/1 | Karos I/1 | 895- mid Xth c. | M2004 | - | N1a1a1a1a4 |
| K1/10 | Karos I/10 | 895- mid Xth c. | CTS1211 | - | R1a1a1b1a2b |
| K1/3286 | Karos I/3286 | 895- mid Xth c. | Z2124 | - | R1a1a1b2a2 |
| K2/6 | Karos II/6 | 895- mid Xth c. | V13 | - | E1b1b1a1b1a |
| K2/33 | Karos II/33 | 895- mid Xth c. | L30;U8 | - | G2a2b |
| K2/26 | Karosc II/26 | 895- mid Xth c. | M253 (xL22) | - | I1 |
| K2/16 | Karos II/16 | 895- mid Xth c. | L621 (xS17250) | - | I2a1a2b |
| K2/52 | Karos II/52 (leader) | 895- mid Xth c. | L621 (xS17250) | - | I2a1a2b |
| K2/51 | Karos II/51 | 895- mid Xth c. | M2004 | - | N1a1a1a1a4 |
| K2/29 | Karos II/29 | 895- mid Xth c. | Z1936 (xL1034) | - | N1a1a1a1a2 |
| K2/61 | Karos II/61 | 895- mid Xth c. | Z2124 | - | R1a1a1b2a2 |
| K2/36 | Karos II/36 | 895- mid Xth c. | Z283 | CTS1211 | R1a1a1b1 |
| K2/41 | Karos II/41 | 895- mid Xth c. | CTS1211 | - | R1a1a1b1a2b |
| K2/18 | Karos II/18 | 895- mid Xth c. | CTS1211 | - | R1a1a1b1a2b |
| K3/12 | Karos III/12 | 895- mid Xth c. | L621 (xS17250) | - | I2a1a2b |
| K3/1 | Karos III/1 | 895- mid Xth c. | M412/U106* | U106 | R1b1a1b1a1a1 |
| K3/13 | Karos III/13 | 895- mid Xth c. | U106 | - | R1b1a1b1a1a1 |
| K3/3 | Karos III/3 | 895- mid Xth c. | M412/U106* | U106 | R1b1a1b1a1a1 |
| KeF1/10936 | Kenézlő-Fazekaszug I/10936 | 895- mid Xth c. | F1096 (xM25) | - | Q1a |
| KeF2/1025 | Kenézlő-Fazekaszug II/1025 | 895- mid Xth c. | M269 (xM412, xU106) | L23 | R1b1a1b |
| KeF2/1027 | Kenézlő-Fazekaszug II/1027 | 895- mid Xth c. | Z1936 (xL1034) | - | N1a1a1a1a2 |
| KeF2/1045 | Kenézlő-Fazekaszug II/1045 | 895- mid Xth c. | Z1936 (xL1034) | - | N1a1a1a1a2 |
| MH/15 | Magyarhomorog/15 | Xth c. | L621 (xS17250) | - | I2a1a2b |
| MH/16 | Magyarhomorog/16 | Xth c. | L621 (xS17250) | - | I2a1a2b |
| MH/9 | Magyarhomorog/9 | Xth c. | M423 (xS17250) | L621 | I2a1a2 |
| SH/41 | Sárrétudvari–Hízó föld/41 | Xth c. second half | U152 | - | R1b1a1b1a1a2b |
| SH/81 | Sárrétudvari–Hízó föld/81 | Xth c. second half | L26 | - | J2a1a |

### East Eurasian Hg-s

Hg Q1a2-M25 is very rare in Europe, where it has highest frequency among Seklers (a Hungarian speaking ethnic group in Transylvania) according to Family Tree DNA database. Ancient samples with Hg Q1a2-M25 are known from the Bronze Age Okunevo and Karasuk cultures, as well as Middle Age Tian Shan Huns and Hunnic-Sarmatians ^18^ implying possible Hunnic origin of this lineage in Europe, which is confirmed by the Hg of our Hun/1 sample, derived from Transylvania. One Conqueror sample KEF1/10936 belongs to Q1a-F1096, possibly to the Q1a1-F746 sister clade of Q1a2-M25, which was not tested in our experiment. Q1a1 is present in East Asia; Mongolia, Japan, China, Korea and its presence in the Kenézlő (KEF) graveyard imply a common substrate of Huns and Conquerors.

Hun/3 belongs to Hg R1a1a1b2a2-Z2124, a subclade of R1a1a1b2-Z93, the east Eurasian subbranch of R1a. Today Z2124 is most frequent in Kyrgyzstan and Afghanistan, but is also widespread among Karachai-Balkars and Baskhirs ^19^. Z2124 was widespread on the Bronze Age steppe, especially in the Afanasievo and Sintashta cultures ^20^ and R1a detected in Xiongnus ^21,22^ very likely belong to the same branch. Two samples from the Karos Conqueror cemeteries (K1/3286 and K2/61) were also classified as R1a-Z2124 and two Avar age individuals (DK/701 and MM/227) belong to the same R1a1a1b2a-Z94 branch but marker Z2124 was not covered in latter samples.

Two Avar samples belonged to Hg C2-M217, which is found mostly in Central Asia and Eastern Siberia. Presence of this Hg had also been detected in the Conquerors, as the Karos2/60 individual belongs to C2 (Fóthi unpublished).

All N-Hg-s identified in the Avars and Conquerors belonged to N1a1a-M178. We have tested 7 subclades of M178; N1a1a2-B187, N1a1a1a2-B211, N1a1a1a1a3-B197, N1a1a1a1a4-M2118, N1a1a1a1a1a-VL29, N1a1a1a1a2-Z1936 and the N1a1a1a1a2a1c1-L1034 subbranch of Z1936. The European subclades VL29 and Z1936 could be excluded in most cases, while the rest of the suclades are prevalent in Siberia ^23^ from where this Hg dispersed in a counter-clockwise migratory route to Europe ^24^. Avar sample MM/58, did not go into any of the tested M178 subclades, while only N1a1a2 could be excluded for the KB/300 Avar khagan due to low coverage. All the 5 other Avar samples belonged to N1a1a1a1a3-B197, which is most prevalent in Chukchi, Buryats, Eskimos, Koryaks and appears among Tuvans and Mongols with lower frequency ^23^. By contrast two Conquerors belonged to N1a1a1a1a4-M2118, the Y lineage of nearly all Yakut males, being also frequent in Evenks, Evens and occurring with lower frequency among Khantys, Mansis and Kazakhs.

### Eurasian Hg-s

Three Conqueror samples belonged to Hg N1a1a1a1a2-Z1936, the Finno-Permic N1a branch, being most frequent among northeastern European Saami, Finns, Karelians, as well as Komis, Volga Tatars and Bashkirs of the Volga-Ural region. Nevertheless this Hg is also present with lower frequency among Karanogays, Siberian Nenets, Khantys, Mansis, Dolgans, Nganasans, and Siberian Tatars ^23^.

### West Eurasian Hg-s

The west Eurasian R1a1a1b1a2b-CTS1211 subclade of R1a is most frequent in Eastern Europe especially among Slavic people. This Hg was detected just in the Conqueror group (K2/18, K2/41 and K1/10). Though CTS1211 was not covered in K2/36 but it may also belong to this sub-branch of Z283.

Hg I2a1a2b-L621 was present in 5 Conqueror samples, and a 6^th^ sample form Magyarhomorog (MH/9) most likely also belongs here, as MH/9 is a likely kin of MH/16 (see below). This Hg of European origin is most prominent in the Balkans and Eastern Europe, especially among Slavic speaking groups. It might have been a major lineage of the Cucuteni-Trypillian culture and it was present in the Baden culture of the Calcholitic Carpathian Basin ^25^.

I1-M253, identified from one Conqueror sample is a northern European Hg found mostly in Scandinavia and Finland and might have originated from this region during the Mesolithic. It has somewhat similar distribution to R1b-U106 associated with Germanic speaking populations.

Three out of 4 samples in the small Karos3 cemetery belonged to Hg R1b1a1b1a1a1-U106 setting apart this cemetery from all other groups, except for the Hun/2 sample which is the only other one with this Hg. Hg U106 is considered a “Germanic” branch as it is most significant today in Germany, Scandinavia, and Britain, and rare in Eastern Europe (Supplementary Table S4). Its ancestral branch Hg R1b1a1b-M262 is assumed to have emerged in the Pontic-Caspian Steppe and arrived to Europe with Bronze Age migrations ^26^. Its presence in Hun and Conqueror samples may derive from Goths, Gepids or other German allies of the Huns.

We detected R1b1a1b1a1a2b-U152 in one sample from the Sárrétudvari Conqueror cemetery, which represent rather commoners than the elite. U152 is the Italo-Celtic R1b branch, concentrated around the Alps, and which was present in the Carpathian Basin before the conquer, so did not necessarily arrived with the Conquerors.

The mediterranean haplogroup E1b1b1a1b1a-V13 was detected in an Avar (SzK/239) and a Conqueror (K2/6) sample, while this marker was not covered in another sample (K1/13, E1b1b-M215). This Hg originated in the Middle East and migrated to the Balkans and Western Asia during the Bronze Age.

The PSZ/1 Avar leader belongs to Hg G2a-P15, while K2/33 to the G2a2b-L30 subbranch. Hg G2a is originated from Anatolia/Iran, now it is most common in the Caucasus region and its is arrival to Europe is associated with the spread of Neolithic farmers ^27^.

One Conqueror sample belongs to Hg J1-M267 and another to J2a1a-L26. Both J1 and J2 lineages are most frequent around the Middle East-Caucasus and probably originated from this region ^28^, then expanded with pastorists prior to the Neolithic.

### Autosomal SNP-s

We have predicted eye, hair and skin color phenotypes from 25 HirisPlex SNP-s, also suitable to predict non-European ancestry ^15^, as summarized in Table 2 and 3. Samples from different archaological cultures and cemeteries showed a remarkable pattern of phenotypic distribution. All Hun and Avar age samples had inherently dark eye/hair colors, DK/701being the only exception (Table 2). Moreover 6/14 Avar age samples were characterized with >0,7 black hair; >0,99 brown eye p-values, inferring 86,5% probability of non-European biogeographic ancestry ^15^ in agreement with their anthropological, archaeological and historical evaluation. In contrast the Conquerors showed a wide variety of phenotypes clustered by cemeteries (Table 3). All individuals from the Sárrétudvari (SH), Magyarhomorog (MH) and majority from the Kenézlő (KEF) graveyards displayed European phenotypic patterns; blue eye and/or light hair with pale skin. In case of Kenézlő this is especially astonishing as all individuals carried east Eurasian mtDNA or Y-chromosomal Hg-s. In the three Karos cemeteries darker eye/hair colors predominated, 4/20 individuals having p-values consistent with non-European origin, nevertheless 5/20 individuals had light hair color indicating a rather mixed origin of this population, concurrent with their mtDNA and Y chromosomal Hg composition.

**Table 2.**
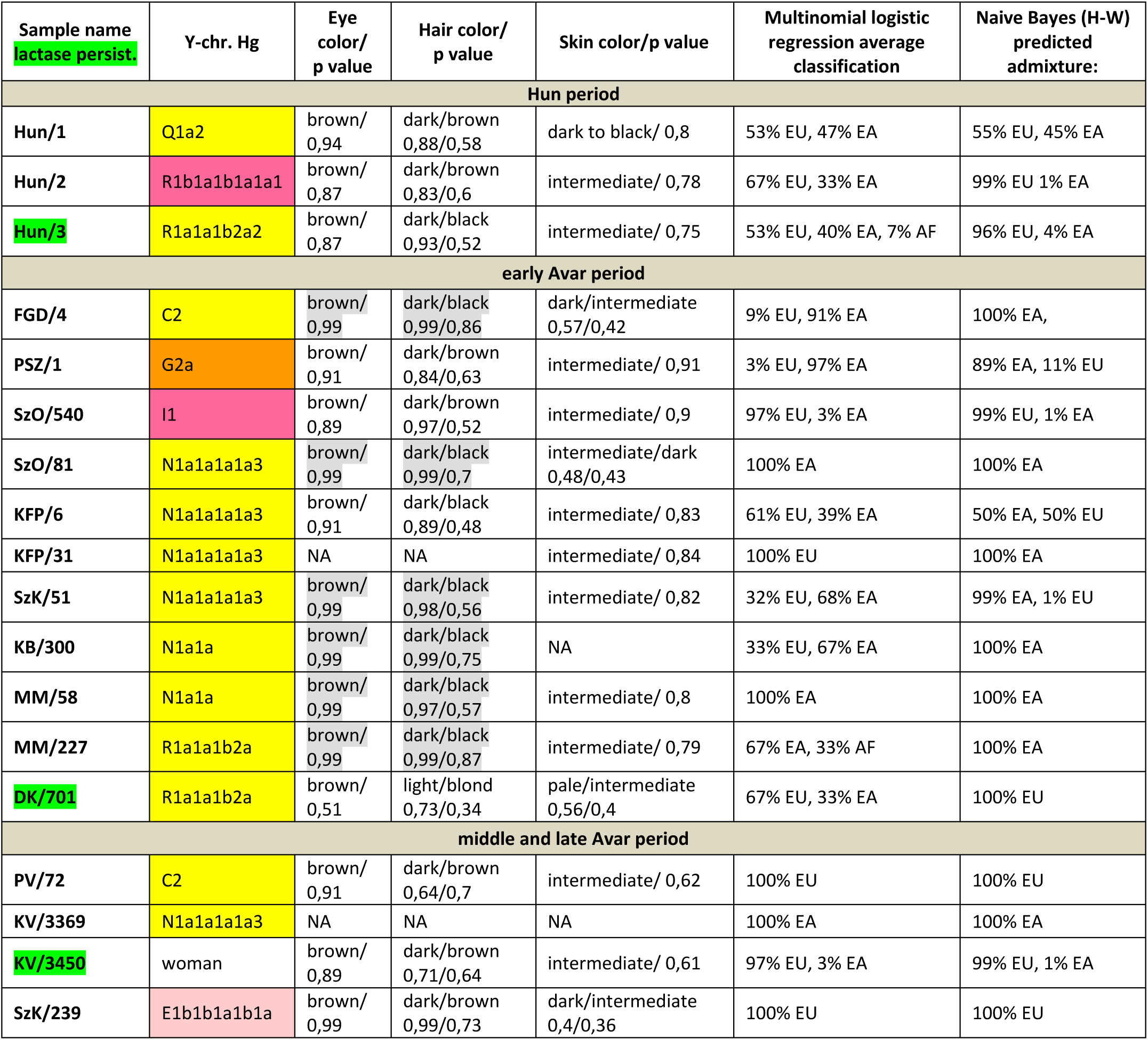
Phenotypes and genetic origin of Hun and Avar age individuals. Eye, hair and skin colors with their probability values were predicted from HIrisPlex markers (Supplementary Table S2), gray shaded values predict non-European ancestry. Most likely origin of individuals were calculated from 34 AIMs (Supplementary Table S2) with two methods; multinomial logistic regression and naive Bayesian classifier (supposing Hardy-Weinberg equilibrium). Names of individuals carrying SNP variants associated with lactase persistence are highlighted with green. Abbreviations are the following: NA=lack of data, EU= Europe, EA=East Asia AF=Africa.

| Sample name<br>lactase persist. | Y-chr. Hg | Eye color/<br>p value | Hair color/<br>p value | Skin color/p value | Multinomial logistic regression average classification | Naive Bayes (H-W) predicted admixture: |
| --- | --- | --- | --- | --- | --- | --- |
| **Hun period** |  |  |  |  |  |  |
| Hun/1 | Q1a2 | brown/<br>0,94 | dark/brown<br>0,88/0,58 | dark to black/ 0,8 | 53% EU, 47% EA | 55% EU, 45% EA |
| Hun/2 | R1b1a1b1a1a1 | brown/<br>0,87 | dark/brown<br>0,83/0,6 | intermediate/ 0,78 | 67% EU, 33% EA | 99% EU 1% EA |
| Hun/3 | R1a1a1b2a2 | brown/<br>0,87 | dark/black<br>0,93/0,52 | intermediate/ 0,75 | 53% EU, 40% EA, 7% AF | 96% EU, 4% EA |
| **early Avar period** |  |  |  |  |  |  |
| FGD/4 | C2 | brown/<br>0,99 | dark/black<br>0,99/0,86 | dark/intermediate<br>0,57/0,42 | 9% EU, 91% EA | 100% EA, |
| PSZ/1 | G2a | brown/<br>0,91 | dark/brown<br>0,84/0,63 | intermediate/ 0,91 | 3% EU, 97% EA | 89% EA, 11% EU |
| SzO/540 | I1 | brown/<br>0,89 | dark/brown<br>0,97/0,52 | intermediate/ 0,9 | 97% EU, 3% EA | 99% EU, 1% EA |
| SzO/81 | N1a1a1a1a3 | brown/<br>0,99 | dark/black<br>0,99/0,7 | intermediate/dark<br>0,48/0,43 | 100% EA | 100% EA |
| KFP/6 | N1a1a1a1a3 | brown/<br>0,91 | dark/black<br>0,89/0,48 | intermediate/ 0,83 | 61% EU, 39% EA | 50% EA, 50% EU |
| KFP/31 | N1a1a1a1a3 | NA | NA | intermediate/ 0,84 | 100% EU | 100% EA |
| SzK/51 | N1a1a1a1a3 | brown/<br>0,99 | dark/black<br>0,98/0,56 | intermediate/ 0,82 | 32% EU, 68% EA | 99% EA, 1% EU |
| KB/300 | N1a1a | brown/<br>0,99 | dark/black<br>0,99/0,75 | NA | 33% EU, 67% EA | 100% EA |
| MM/58 | N1a1a | brown/<br>0,99 | dark/black<br>0,97/0,57 | intermediate/ 0,8 | 100% EA | 100% EA |
| MM/227 | R1a1a1b2a | brown/<br>0,99 | dark/black<br>0,99/0,87 | intermediate/ 0,79 | 67% EA, 33% AF | 100% EA |
| DK/701 | R1a1a1b2a | brown/<br>0,51 | light/blond<br>0,73/0,34 | pale/intermediate<br>0,56/0,4 | 67% EU, 33% EA | 100% EU |
| **middle and late Avar period** |  |  |  |  |  |  |
| PV/72 | C2 | brown/<br>0,91 | dark/brown<br>0,64/0,7 | intermediate/ 0,62 | 100% EU | 100% EU |
| KV/3369 | N1a1a1a1a3 | NA | NA | NA | 100% EA | 100% EA |
| KV/3450 | woman | brown/<br>0,89 | dark/brown<br>0,71/0,64 | intermediate/ 0,61 | 97% EU, 3% EA | 99% EU, 1% EA |
| SzK/239 | E1b1b1a1b1a | brown/<br>0,99 | dark/brown<br>0,99/0,73 | dark/intermediate<br>0,4/0,36 | 100% EU | 100% EU |

**Table 3.** Phenotypes and genetic origin of Conqueror period individuals. mtDNA Hg-s were taken from ^11^, all markings are identical to that of Table 1.

| Sample name<br>lactase persist. | Y-chr. Hg | mtDNA Hg | Eye color/p value | Hair color/p value | Skin color/p value | Multinomial logistic regression average classification | Naive Bayes (H-W) predicted admixture: |
| --- | --- | --- | --- | --- | --- | --- | --- |
| **Conqueror period** |  |  |  |  |  |  |  |
| K1/13 | E1b1b | U5b2b3 | NA | NA | NA | 100% EU | 99% EU, 1% EA |
| K1/1438 | J1 | U3b1b | NA | NA | NA | 100% EU | 100% EU |
| K1/1 | N1a1a1a1a4 | H1b2 | brown/ 0,91 | dark/brown 0,59/0,51 | intermediate/ 0,99 | 35% EU, 65% EA | 97% EU, 3% EA |
| K1/10 | R1a1a1b1a2b | U3b1b | brown/ 0,63 | dark/brown 0,7/0,64 | intermediate/pale 0,58/0,39 | 100% EU | 100% EU |
| K1/3286 | R1a1a1b2a2 | T2b4h | brown/ 0,91 | dark/black 0,88/0,56 | intermediate/ 0,99 | 100% EA | 76% EA, 24% EU |
| K2/6 | E1b1b1a1b1a | H5v | blue/ 0,95 | light/red 0,9/0,51 | NA | 100% EU | 100% EU |
| K2/33 | G2a2b | U4a1b1 | brown/ 0,99 | dark/brown 0,73/0,54 | intermediate/ 0,69 | 100% EU | 100% EU |
| K2/26 | I1 | T1a1 | brown/ 0,99 | dark/black 0,99/0,76 | intermediate/ 0,57 | 33% EU, 67% EA | 60% EA, 40% EU |
| K2/16 | I2a1a2b | X2I | brown/ 0,98 | dark/brown 0,81/0,72 | intermediate/ 0,58 | 52% EA, 13% EU, 35% AF | 98% EU, 2% EA |
| K2/52 | I2a1a2b | X2f | brown/ 0,71 | light/brown 0,64/0,58 | intermediate/ 0,57 | 2% EU, 98% EA | 100% EU |
| K2/51 | N1a1a1a1a4 | U4d2 | brown/ 0,99 | dark/black 0,99/0,86 | dark/ 0,641 | 100% EU | 100% EU |
| K2/29 | N1a1a1a1a2 | J1b1a1e | brown/ 0,77 | light/blond 0,85/0,5 | intermediate/pale 0,5/0,48 | 37% EU, 30% EA, 33% AF | 54% EA, 46% EU |
| K2/61 | R1a1a1b2a2 | U4d2 | brown/ 0,99 | dark/black 0,98/0,73 | intermediate/ 0,98 | 100% EA | 100% EA |
| K2/36 | R1a1a1b1 | D4i2 | brown/ 0,63 | dark/brown 0,72/0,7 | intermediate/ 0,52 | 94% EU, 6% EA | 100% EU |
| K2/41 | R1a1a1b1a2b | H35 | NA | NA | NA | 100% EA | 100% EA |
| K2/18 | R1a1a1b1a2b | T1a1 | brown/ 0,84 | light/brown 0,6/0,6 | intermediate/ 0,69 | 50% EU, 50% EA | 85% EU, 15% EA |
| K3/12 | I2a1a2b | A12 | brown/ 0,99 | dark/black 0,98/0,53 | dark/ 0,68 | 7% EU, 93% EA | 100% EU |
| K3/1 | R1b1a1b1a1a1 | B4d1 | brown/ 0,74 | dark/ 0,76 | NA | 67% EU, 33% EA | 93% EU, 7% EA |
| K3/13 | R1b1a1b1a1a1 | B4d1 | brown/ 0,84 | light/brown 0,71/0,46 | intermediate/ 0,99 | 67% EA, 33% EA | 100% EA |
| K3/3 | R1b1a1b1a1a1 | H6a1b | NA | NA | intermediate/ 0,55 | 100% EU | 100% EU |
| K3/6 | woman | B4d1 | brown/ 0,98 | dark/brown 0,8/0,66 | intermediate/ 0,77 | 35% EU, 1% EA, 64% AF | 99% EA, 1% EU |
| KEF1/10936 | Q1a | H6a1b | brown/ 0,67 | light/red 1,0/0,64 | intermediate/pale 0,56/0,33 | 38% EU, 62% EA | 96% EA, 4% EU |
| KEF2/1025 | R1b1a1b | G2a1 | brown/<br>0,97 | dark/black<br>0,97/0,55 | intermediate/ 0,97 | 100% EA | 100% EA |
| KEF2/1027 | N1a1a1a1a2 | N1a1a1a1a | blue/<br>0,69 | dark/brown<br>0,97/0,53 | intermediate/ 0,62 | 100% EU | 100% EU |
| KEF2/1045 | N1a1a1a1a2 | N1a1a1a1a | blue/<br>0,64 | light/brown<br>1,0/0,51 | very pale/ 0,54 | 100% EU | 100% EU |
| MH/15 | I2a1a2b | NA | brown/<br>0,48 | dark/black<br>0,91/0,68 | pale/ 0,63 | 67% EU, 33% EA | 100% EU |
| MH/16 | I2a1a2b | NA | blue/<br>0,91 | light/blond<br>0,96/0,37 | intermediate/pale<br>0,57/0,41 | 67% EA, 33% EA | 100% EU |
| MH/9 | I2a1a2 | NA | blue/<br>0,95 | light/ 0,95 | pale/ 0,59 | 67% EU, 33% EA | 100% EU |
| SH/41 | R1b1a1b1a1a2b | NA | brown/<br>0,59 | light/brown<br>0,66/0,48 | pale/intermediate<br>0,5/0,45 | 72% EU, 26% EA,<br>2% AF | 96% EU, 4%<br>EA |
| SH/81 | J2a1a | H1c | brown/<br>0,67 | light/brown<br>0,79/0,42 | intermediate/pale<br>0,54/0,43 | 67% EU, 34% EA | 100% EU |
| SH/103 | woman | C4a1b | brown/<br>0,77 | light/blond<br>0,93/0,44 | intermediate/ 0,97 | 32% EU, 37% EA,<br>31% AF | 93% EA, 7%<br>EU |

We have also determined 34 AIMs, which can inform about the likely geographic origin of individuals, and used the Snipper App suite version 2.5 portal ^17^ to assign biogeographic ancestry. We predicted ancestry with both the naive Bayesian classifier and multinomial logistic regression (MLR) algorithms, as these make different assumptions about genetic equilibrium ^29^, and listed the results on Tables 2 and 3. The AIM-s results fairly matched and complemented phenotypic information. All Hun age individuals revealed admixture derived from European and East Asian ancestors, while 8/15 Avar age individuals showed predominantly East Asian origin with both methods, 4 individuals were definitely European, while two showed evidence of admixture. The KFP/31 sample gave contradicting results due to low coverage.

Conqueror samples from the Magyarhomorog (MH) and Sárrétudvari (SH) cemeteries showed mostly European ancestry in agreement with their phenotypes and Y Hg-s, though MLR detected a significant east Asian ancestry component and the SH/103 women was classified east Asian despite her blond hair. The Karos (K) and Kenézlő (KEF) populations were profoundly admixed, comprising individuals of purely East Asian, European and mixed origin in nearly identical proportions, again in agreement with results obtained from uniparental and phenotypic markers.

The determined variable autosomal loci are also suitable to exclude possible direct (parent-child, sibling) genetic relatedness ^30^, thus we compared the autosomal genotypes of all individuals sharing either maternal or paternal Hg-s. Direct kinship could not be excluded between the MH/9 and MH/16 individuals with identical phenotypes, suggesting that MH/9 probably also belongs to Hg I2a1a2b-L621, despite its uncovered L621 marker. KEF2per1027 and KEF2per1045 were probably brothers as they had identical mitogenomes, Y Hg-s and blue eye color besides sharing autosomal alleles. The same applies to K3/1 and K3/13 individuals who were probably also brothers. We could not exclude possible direct paternal relationship between K2/36, K2/18 and K2/41 but the first two samples had unsatisfactory coverage to make a strong statement.

We have tested two SNP-s (rs4988235 and rs182549) associated with the adult lactase persistence phenotype in Europe ^16^. Individuals carrying derived alleles in these loci are able to digest lactose in dairy products during adulthood without symptoms of lactose intolerance. Allele frequency of the persistence genotype varies throughout Eurasia ^31^, reaching above 90% at some parts of Northwestern Europe, around 80% in present day Hungary, but drops below 30% in Central Asia and even lower in East Asia. We detected the derived persistence allele in all of the studied groups (Table 2 and 3, Supplementary Table S2); 1/3 of the Hun period individuals, 2/14 of the Avar period individuals, and 6/31 of the Conqueror period individuals carried persistence alleles. It is remarkable that the persistence genotype seems to be strongly associated with European origin, as all of the carriers were predicted to have predominantly European ancestors. This is in agrrement with a previous study ^32^, which found that all of the 11% Conqueror samples with persistence genotype carried European mtDNA Hg H. In addition all carriers were heterozygous, 6 of them for both SNP-s, but three of them carried just the rs182549 (−22018 G>A) derived allele suggesting previous admixture with non-carriers, possibly derived from East Eurasia.

### Population genetic analysis

The studied Conqueror group very likely represent real populations, as 24 of the 29 samples came from 4 nearby cemeteries (Karos 1,2,3 and Kenézlő) with identical archaeological and anthropological groupings and the studied Magyarhomorog individuals were also categorized as belonging to the same early Conqueror elite. The two Sárrétudvari samples however represent commoners from the second half of the 10^th^ century and their Hg-s also reveal this difference, thus they were excluded from the population genetic analysis (though including them does not alter the results). The Avar group was assembled from several different cemeteries of a wider timespan, thus they cannot represent the Avar period population of the Carpathian Basin, however their relatively homogenous Hg distribution indicate that the Avar elite embodied the same east Eurasian sub-population throughout their reign, so it appeared to be meaningful to include them in the analysis.

In order to find the most similar populations to our studied ones, we compared the Hg distribution of the Avar and Conqueror elite to that of 78 modern Eurasian populations (Supplementary Table S4) and represented their relations on MDS plot displayed on Fig. 2. In order to increase resolution, we separately calculated the frequencies of each sub-Hg-s which are present in our samples, while frequencies of all other subclades were combined as listed in Supplementary Table S4.

**Fig. 2.**
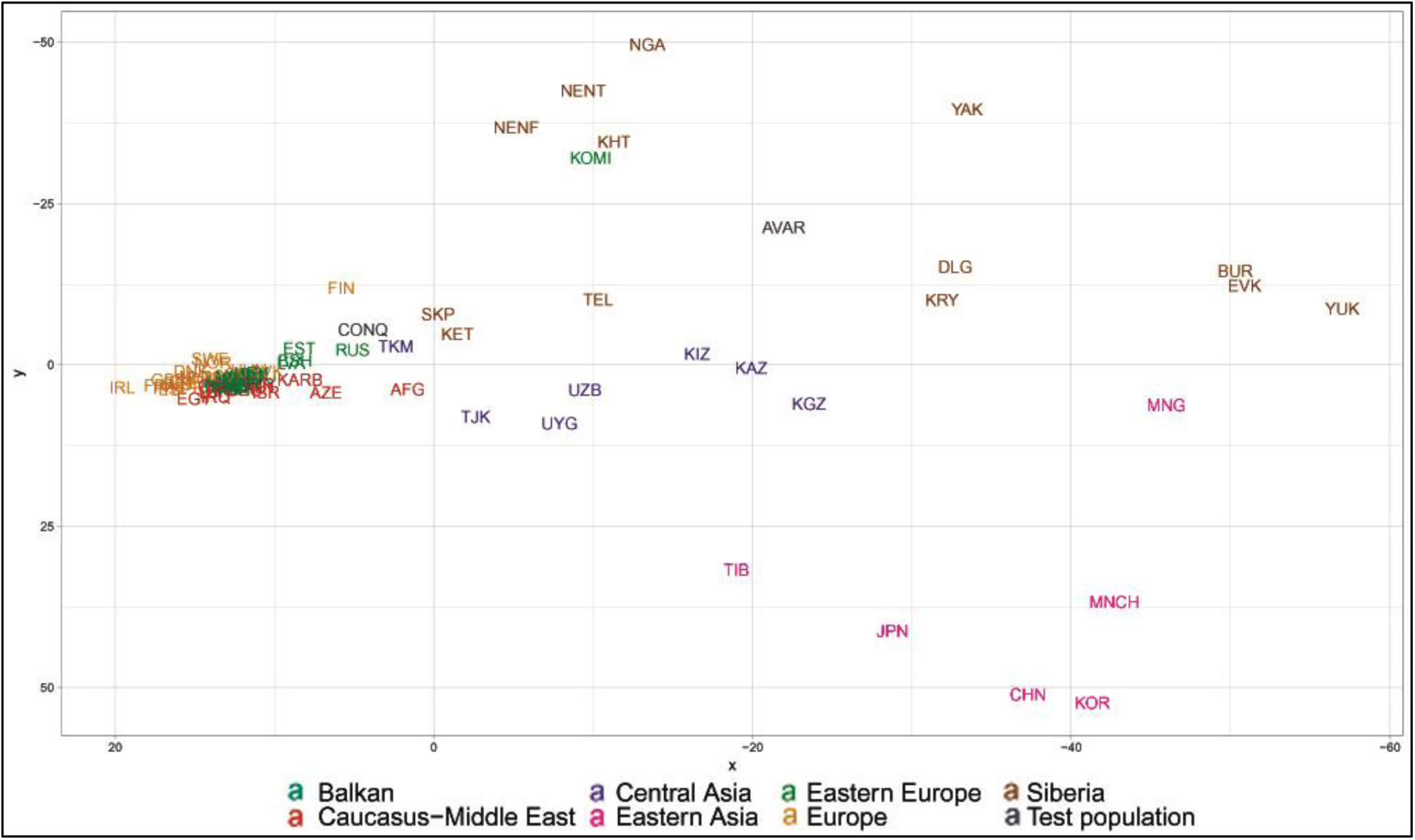
MDS plot of Y-chromosomal Hg distribution of 78 Eurasian populations including Avars and Conquerors. Population three letter codes are given in Supplementary Table S4.

Similar Hg distributions are mapped into neighboring positions on the MDS plot, which clearly separates populations according to their geographic locations. Along the *x* axis east Eurasian populations map to the right while Europeans are compressed at the left. Along the *y* axis Siberian populations are sequestered at the top, while East Asian ones at the bottom of the graph. The Avars are obviously mapped to the Siberian side with smallest weighted Euclidean distances from Koryaks (KRY), Teleuts (TEL), Khantys (KHT), Komis, (KOM) and Dolgans (DLG). The Conquerors are positioned between eastern Europeans, Central Asians and Siberians but their exact relations are hard to make out because of the crowding at the European side.

In order to better discern the position of the Conquerors we redraw the MDS plot without the eastern Asian and Siberian populations (Fig. 3). Though having removed subset of the data MDS imaging in two-dimensions inevitably rearranged the remaining components, but the general organization of the population clusters remained unchanged: Western-Northern Europeans map to the left, Central-eastern European and Balkan people around the middle, Central Asians to the top right, and Caucasus-Middle East populations to bottom right. Conquerors remained between Central Asians and eastern Europeans, with smallest weighted Euclidean distances from Bashkirs (BSH), Hungarians (HUN), Tajiks (TJK), Estonians (EST), Kazakhs (KAZ), Uzbeks (UZB) and Slovaks (SVK).

**Fig. 3.**
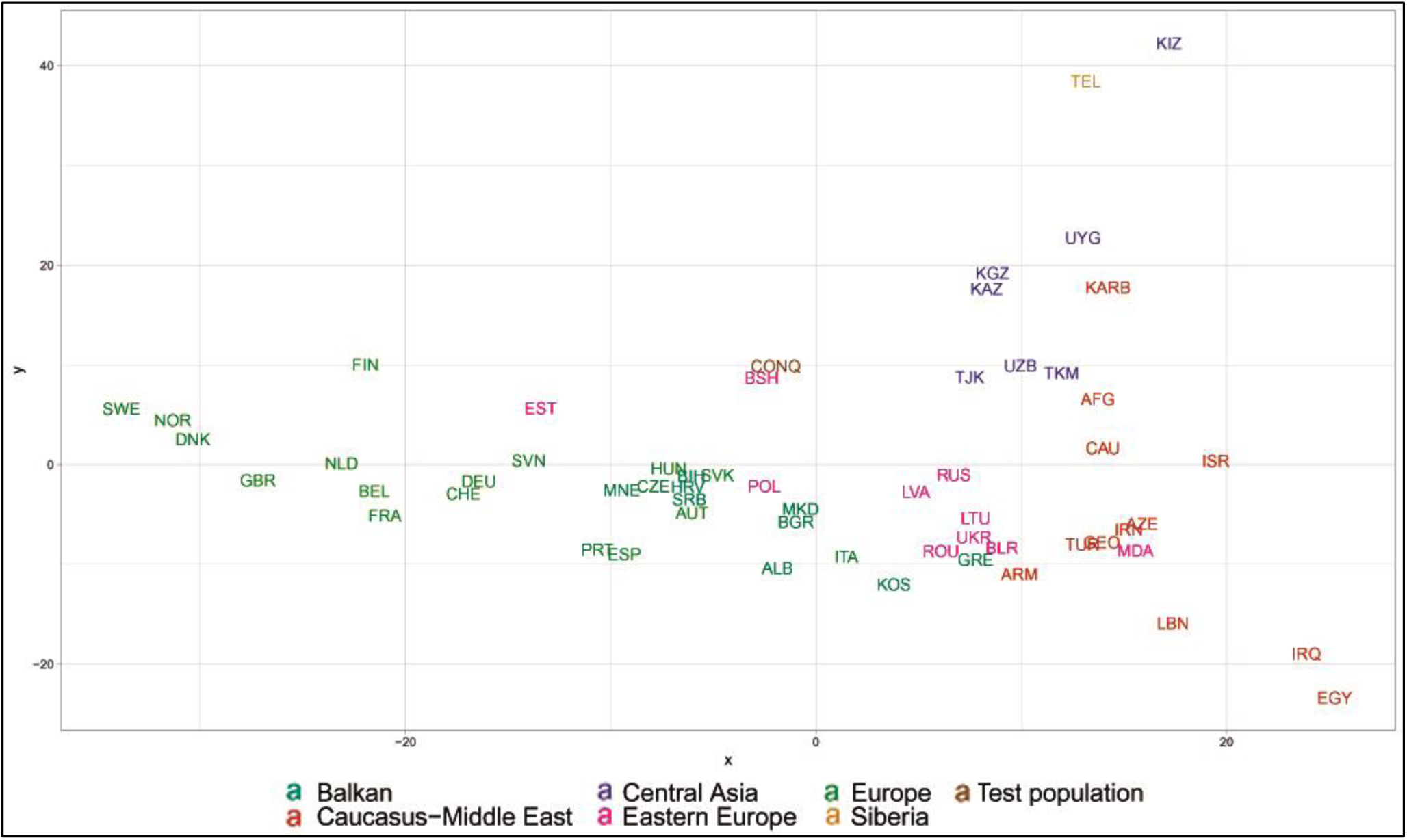
MDS plot of Y-chromosomal Hg distribution of 58 European and Central Asian populations including Conquerors. Population three letter codes are given in Supplementary Table S4.

The considerable frequency of Hg N1a in Conquerors and especially in Avars facilitates another analysis in which the frequency distribution of their N1a subbranches can be compared to that of all Eurasian populations carrying this Hg, which is described in ^23^. The N1a database of ^23^ contains several relevant Eurasian populations missing from Supplementary Table S3, moreover the Y dataset of Supplementary Table S4 has a low resolution of N1a subclades, thus this analysis is expected to provide additional information. We combined the N1a dataset published in ^23^ with our data as presented in Supplementary Table S5 and performed Principal Component Analysis (PCA) as shown on Fig. 4.

**Fig. 4.**
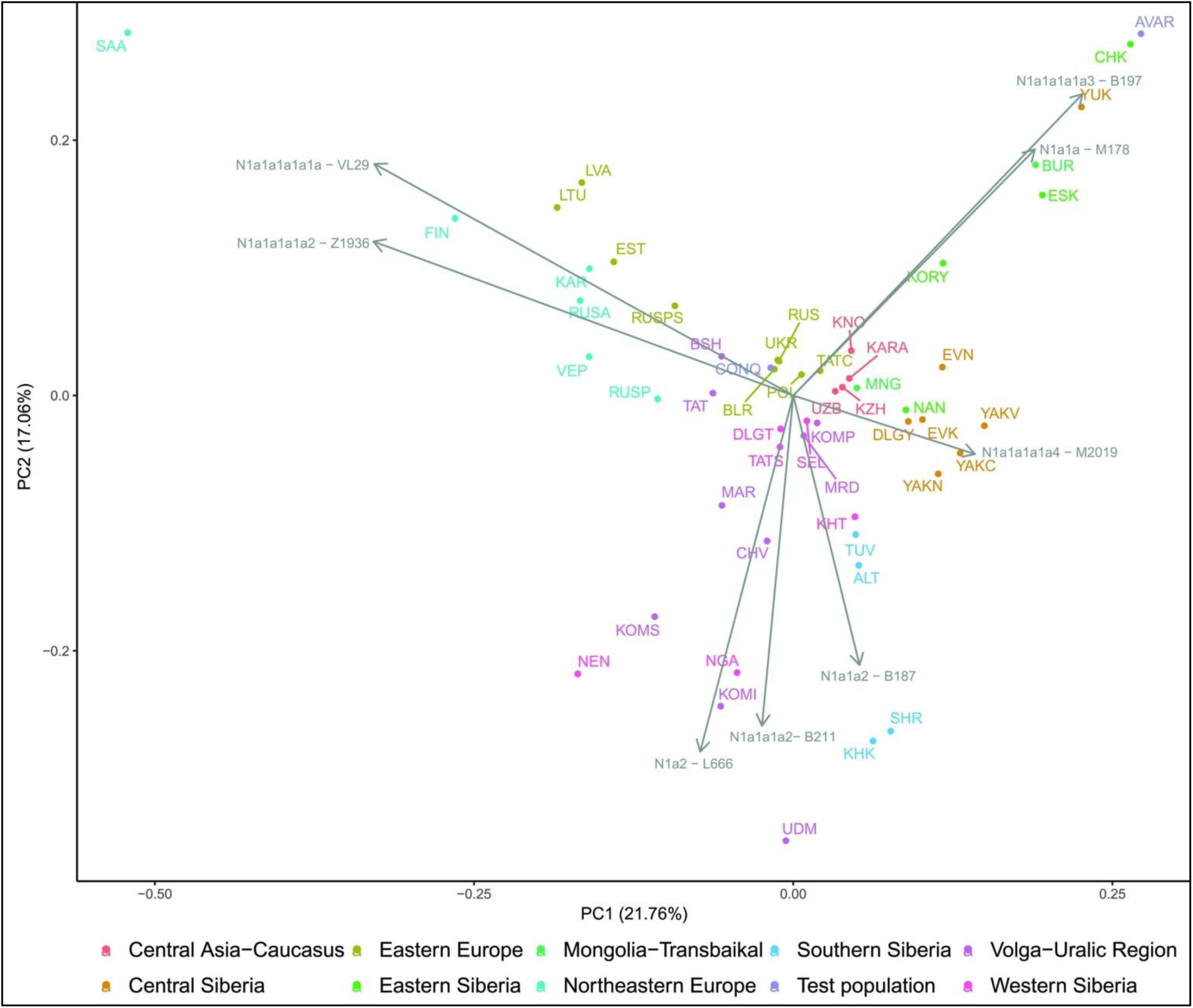
First two components of PCA from Hg N1a subbranch distribution in 51 populations including Avars and Conquerors. Colors indicate geographic regions. Three letter codes are given in Supplementary Table S5.

PCA also separates populations according to their geographical positions, as identical colors appear in the same segments of the graph. The Avars map to the top right corner, close to eastern Siberian Chukchis (CHK), Yukaghirs (YUK), Eskimos (ESK) and Transbaikalian Buryats (BUR). The Conquerors (CONQ) map to the middle of the graph, very close to eastern European Belorusians (BLR), Ukrainians (UKR), Russians (RUS) and Polish (POL), not far from Volga-Uralic region Bashkirs (BSH), Volga Tatars (TAT), Komi Permians (KOMP) and Mordvins (MRD), also surrounded by Central Asian groups; Karanogays (KNO), Karakalpaks (KARA), Uzbeks (UZB) as well as Crimean Tatars (TATC), Mongols (MON), Selkups (SEL) and Dolgans from Taymyr (DLGT).

## Discussion

The origin and composition of the Conqueror paternal lineages fairly mirrors that of their maternal ones^11^; 20,7% of the Y-Hg-s originated from East Eurasia, this value is 30,4% for mtDNA; proportion of west Eurasian paternal lineages is 69% compared to 58,8% for mtDNA; while proportion of lineages with north-western European and Caucasus-Middle East origin are nearly the same affirming that both males and females of similar origin migrated together. Both MDS analysis of the entire Conqueror Y chromosome pool and PCA of their N1a lineages indicates that their admixture sources are found among Central Asians and eastern European Pontic Steppe groups, a finding comparable to what had been described for maternal lineages ^11^. Composition of the Conqueror paternal lineages is very similar to that of Baskhirs, while their maternal composition was found most similar to Volga Tatars ^11^. These modern populations are located next to each other, have similar prehistory _33_ and genetic structure derived from the same admixture sources detected in the Conquerors. Moreover it must be noted that Bashkirs were not represented in the mitogenome database while Volga Tatars are missing from our Y-chromosome database due to lack of data, but their N1a distribution is quite similar (Fig. 4), thus mtDNA results are in accord with Y-chromosomal ones.

The Conqueror-Bashkir relations are also supported by historical sources, as early Hungarians of the Carpathian Basin were reported to be identical to Baskhirs by Arabic historians like al-Masudi, al-Qazwini, al-Balhi, al-Istahri and Abu Hamid al-Garnati ^34^, latter visited both groups at the same time around 1150 AD and used the term Bashgird to refer to the Hungarians in the Carpathian Basin. In addition parallels were found between several Conqueror and Bashkir tribe names and Bashkiria has been identified with Magna Hungaria, the motherland of Conquerors ^35^.

We identified potential relatives within Conqueror cemeteries but not between them. The uniform paternal lineages of the small Karos3 (19 graves) and Magyarhomorog (17 graves) cemeteries approve patrilinear organization of these communities. The identical I2a1a2b Hg-s of Magyarhomorog individuals appears to be frequent among high-ranking Conquerors, as the most distinguished graves in the Karos2 and 3 cemeteries also belong to this lineage. The Karos2 and Karos3 leaders were brothers with identical mitogenomes ^11^ and Y-chromosomal STR profiles (Fóthi unpublished). The Sárrétudvari commoner cemetery seems distinct from the others, containing other sorts of European Hg-s. Available Y-chromosomal and mtDNA data ^11^ from this cemetery suggest that common people of the 10^th^ century rather represented resident population than newcomers. The great diversity of Y Hg-s, mtDNA Hg-s, phenotypes and predicted biogeographic classifications of the Conquerors indicate that they were relatively recently associated from very diverse populations.

In contrast the studied Avar military leader group had a much more uniform origin. The Avar group carried predominantly East Eurasian lineages in accordance with their known Inner Asian origin inferred from archaeological and anthropological parallels as well as historical sources. However the unanticipated prevalence of their Siberian N1a Hg-s, sheds new light on their prehistory. Accepting their presumed Rouran origin would implicate a ruling class with Siberian ancestry in Inner Asia before Turkic take-over. The surprisingly high frequency of N1a1a1a1a3 Hg reveals that ancestors of contemporary eastern Siberians and Buryats could give a considerable part the Rouran and Avar elite, nevertheless a larger sample size from more Avar cemeteries are needed to clarify their exact composition.

The genetic profile of the Avar and Conqueror leader groups seems considerably different, as latter group is distinguished by the significant presence of European Hg-s; I2a1a2b-L621, R1b1a1b1a1a1-U106 and the Finno-Permic N1a1a1a1a2-Z1936 branch. Their Siberian N1a1a1a1a4 subclade also points at different source populations among ancestors of Yakuts, Evenks and Evens. Nevertheless the east Eurasian R1a subclade, R1a1a1b2a-Z94 seems to be a common element of the Hun, Avar and Conqueror elite. In contrast to Avars, all three Hun lineages have paralleles among the Conquerors, but strong inferences cannot be drawn due to small sample size.

It is generally accepted that the Hungarian language was brought to the Carpathian Basin by the Conquerors. Uralic speaking populations are characterized by a high frequency of Y-Hg N, which have often been interpreted as a genetic signal of shared ancestry. Indeed, recently a distinct shared ancestry component of likely Siberian origin was identified at the genomic level in these populations, modern Hungarians being a puzzling exception^36^. The Conqueror elite had a significant proportion of N Hgs, 7% of them carrying N1a1a1a1a4-M2118 and 10% N1a1a1a1a2-Z1936, both of which are present in Ugric speaking Khantys and Mansis ^23^. At the same time none of the examined Conquerors belonged to the L1034 subclade of Z1936, while all of the Khanty Z1936 lineages reported in ^37^ proved to be L1034 which has not been tested in the ^23^ study. Population genetic data rather position the Conqueror elite among Turkic groups, Bashkirs and Volga Tatars, in agreement with contemporary historical accounts which denominated the Conquerors as “Turks”^38^. This does not exclude the possibility that the Hungarian language could also have been present in the obviously very heterogeneous, probably multiethnic Conqueror tribal alliance.

## Methods

All libraries were made from partial uracil-DNA-glycosylase (UDG) treated DNA extracts. Details of the aDNA extraction, NGS library construction, sequencing and sequence analysis methods are given in ^11,39^. For designing oligonucleotide probes we selected 120 nucleotide (nt) long sequences around each targeted SNP (listed in Supplementary Table S1 and S2), with the SNP in central position, and designed 80 nt long oligos with 3x tiling and 60 nt overlap. Selected sequences were sent to MYcroarray (Ann Arbor, MI, USA, now Arbor Biosciences) for quality control. Sequences with acceptable quality were used to synthetize a custom biotinylated MYbaits RNA oligonucleotide probe set at MYcroarray.

We tested the endogenous DNA content of each library with low coverage shotgun sequencing (Supplementary Table S6) and omitted samples with low human DNA content. Enrichment was done from libraries containing filled-in short (33 and 34 nt long) adapters; IS1_adapter_P5 and IS2_adapter_P7 ^40^. These libraries were preamplified in 2 x 50 µl reactions containing 800 nM each of IS7_short_amp.P5 and IS8_short_amp.P7 primers, 200 µM dNTP mix, 2 mM MgCl_2_, 0,02 U/µl GoTaq G2 Hot Start Polymerase (Promega) and 1X GoTaq buffer, followed by MinElute purification. PCR conditions were 96 ^o^C 6 min, 16 cycles of 94 ^o^C 30 sec, 58 ^o^C 30 sec, 72 ^o^C 30 sec, followed by a final extension of 64 ^o^C 10 min. Libraries were eluted from the column in 50 µl 55 ^o^C EB buffer (Qiagen), and concentration was measured with Qubit (Termo Fisher Scientific). Then 30-50 ng preamplified library was amplified again in the same reaction with 7-8 cycles to obtain enough DNA for enrichment.

Libraries were enriched individually without pooling according to the instructions provided by the manufacturer (MYbaits manual v3.0) with the following specifications: Just one round of enrichment was done, the enrichment reaction contained 300 ng preamplified library and 100 ng RNA probe, supplemented with 33,3 ng/ul yeast tRNA carrier. The following short phosphorylated blocking oligos were used to mask adapters in 27,3 ng/μl concentrations: BO2.P5.part2F: 5’ ACACTCTTTCCCTACACGACGCTCTTCCGATCT[Phos] 3’ BO4.P7.part1R: 5’ GTGACTGGAGTTCAGACGTGTGCTCTTCCGATCT[Phos] 3’ Hybridization was done at 65 ^o^C for 72 hours, then bead-bound enriched libraries were resuspended in 24 μl water and released from the beads in a 50 μl PCR reaction containing 1 X KAPA HiFi HotStart ReadyMix and 1000 nM each of P5 forward and P7 reverse indexing primers. PCR conditions were: 98 ^o^C 2 min, 16 cycles of 98 ^o^C 20 sec, 60 ^o^C 30 sec, 72 ^o^C 30 sec, followed by a final extension of 72 ^o^C 1 min. The captured and amplified library was purified on MinElute column and eluted in 16 μl EB. Quantity and quality measurements were performed using Qubit and TapeStation (Agilent).

Sequencing was done on Illumina MiSeq platform (2x 150 bp paired end reads) or NextSeq platform (1x 150 bp single end reads). Adapters were trimmed with the cutadapt software ^41^ in paired or single end read mode. Read quality was assessed with FastQC ^42^. Sequences shorter than 25 nucleotide were removed from the dataset, the resulting reads were mapped to the GRCh37.75 human genome reference sequence using the Burrows Wheeler Aligner (BWA) v0.7.9 software ^43^ with the BWA mem algorithm and default parameters. In case the read counts were insufficient, according to duplicate statistics and library complexity, new library preparation and/or new NGS sequencing were performed and the aligned BAM files from multiple runs were merged by the samtools (version 1.3.1) merge algorithm ^44^. Duplicates were removed by PICARD tools (version 1.113) ^45^ MarkDuplicates algorithm and < MAPQ 50 reads were excluded from downstream analysis. The SNPs were genotyped by freebayes (version 1.2.0) [http://arxiv.org/abs/1207.3907] using the „–variant-input” parameter with a VCF file and the „-t” flag with a bed file containing the coordinates of the SNPs. For each samples the reference and alternate allele counts for each positions were exported to a joint XLS file, and variants with low allele counts were also manually evaluated in IGV.

Five N1a Y-chromosomal sub-Hg-s were determined by amplicon sequencing. Annealing temperatures were optimized for each primers and non-template PCR controls were done for each reaction. PCR fragments were purified from agarose gels and sequenced on Ion Torrent Personal Genome Machine (PGM). Primers with amplified Hg defining markers, annealing temperatures and product length are listed below: N1a1a1a1a3 (B197) GRCh37: 14827819 T>C, product Size: 98 bp, annealing temp: 58,7 ^o^C B197_F: GGATGCACTGATAATCCTTAACTTT, B197_R: TTGACTAAGAAAAAGAGAAGAGTCAAA N1a1a1a1a3 (Z35291) GRCh37: 16510372 C>A, product Size: 82 bp, annealing temp: 57,3 ^o^C Z35291_F: AACAAGACATACATAGAAGAGCTGTG, Z35291_R: CGAAGAGAAATGGATGATTCG Y24374_F: TGTCCAATCACTGTCCATGC, Y24374_R: TGAAACTTTCTATCTGTTCATCTGC N1a1a1a1a4 (M2004): GRCh37: 7602098 T>C, product Size: 60 bp, annealing temp: : 56.2 ^o^C YM2004_F: TTTTCAAAGGCTTGGTAGAGG, YM2004_R: AAAAAGACTACTCATCCTTTCTGTTT N1a1a1a2 (Y9286): GRCh37: 16220297 C>T, product Size: 81 bp, annealing temp: : 56.2 ^o^C Y9286_F: TGCAGCTTAAGAACTCATGAAGA, Y9286_R: AAATCTGCTCACTCTCTATTTGACT N1a1a2 (Z35159): GRCh37: 2869260 T>C, product Size: 60 bp, annealing temp: 56.2 ^o^C YZ35159_F: ACCTAAAAACACTACCAACAGAGTG, YZ35159_R: TGACAGATATTTTCCCTATTTGTGG PCR conditions were the following: 1 μl preamplified library DNA, 1.5 mM MgCl_2_, 0.4 μM of each primers, 0.1 mM of each dNTPs, 0.6 mg/ml BSA and 1 U of GoTaqG2 Hot Start Polymerase, 1x Green GoTaq Buffer. Thermocycling conditions were 96 ^o^C 6 min, followed by 45 cycles of 95 ^o^C 30 sec, annealing temp. 30 sec, 72 ^o^C 30 sec, and final extension at 65 ^o^C for 5 min. Determined amplicon sequences were integrated into the original BAM files.

We used the HIrisPlex-S eye, hair and skin color DNA phenotyping webtool https://hirisplex.erasmusmc.nl/ ^15^ for phenotype determination and the Snipper App suite version 2.5 portal http://mathgene.usc.es/snipper/ ^17^ to assign biogeographic ancestry.

Y-Hg population database of 77567 individuals from 79 modern Eurasian populations was assembled from FamilyTreeDNA Y-DNA Haplotree https://www.familytreedna.com/public/y-dna-haplotree/A or FamilyTreeDNA Geographical Projects https://www.familytreedna.com/projects.aspx supplemented with data from ^37^ (Supplementary Table S4). The Eurasian N1a Hg database was taken from ^23^ (Supplementary Table S5). Classical (metric) MDS was performed using the cmdscale function in MASS package of R ^46^, while PCA was done with the prcomp function of R.

## Data availability

BAM files were deposited to the European Nucleotide Archive under accession no: PRJEB31764

## Acknowledgements

We would like to thank György Szabados for his useful advices in historical background. László Révész, László Kovács and Balázs Tihanyi helped us in topics of archaeology. We are grateful to Erzsébet Fóthi for sharing unpublished results. We thank Horolma Pamjav for her help in marker selection, and Péter Bihari for his help with next generation sequencing.

This work was supported by grants from the National Research, Development and Innovation Office (K-124350 to TT) and The House of Árpád Programme (2018-2023) Scientific Subproject: V.1. Anthropological-Genetic portrayal of Hungarians in the Arpadian Age to TT.

## Author Contributions

Conceptualization: TT, EN

Data curation: EN, ZM

Formal analysis: EN, ZM, TT

Funding acquisition: TT

Investigation: EN, KM, DL,

Methodology: EN, ZM, IN, TT

Project administration: TT

Resources: ÁK, EM, AM, GYP, CSB, PT, SZG, BK, LK

Software: ZM

Supervision: TT

Visualization: EN, TT

Writing – original draft: TT

Writing – review & editing: EN, ZM, TK, IR, IN, CSB

## Additional Information

Dr. Latinovics and Dr. Nagy had consulting positions at SeqOmics Biotechnology Ltd. at the time the study was conceived. SeqOmics Biotechnology Ltd. was not directly involved in the design and execution of the experiments or in the writing of the manuscript.

## Supplementary Information

Archaeological and anthropological background

**Supplementary Table S1.** Y SNP list, and Y chromosomal data

**Supplementary Table S2.** Autosomal SNP list and data

**Supplementary Table S3.** NGS data

**Supplementary Table S4** Y chromosomal population database

**Supplementary Table S5.** N1a population database

**Supplementary Table S6.** Shotgun sequence data

