## Supplementary Information for "Y-chromosome haplogroups from Hun, Avar and conquering Hungarian period nomadic people of the Carpathian Basin"

### Archaeological and anthropological background:

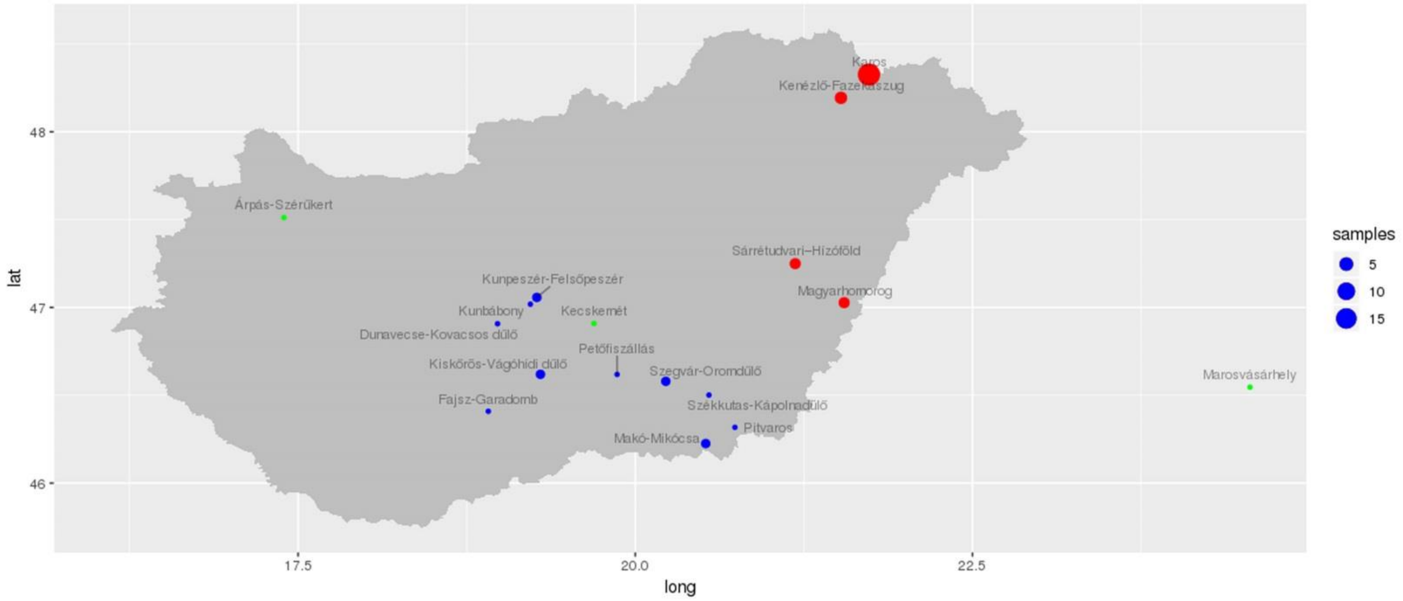

**Location of the cemeteries** reported in this study**.** Red dots indicate Conqueror cemeteries, blue dots Avar age cemeteries, green dots Hun age cemeteries. Dot size is proportional to sample size as indicated. Map was created with the maps package of R [24].

***Hun age graves:***

**Árpás-Szérűskert-1**

The lonely grave of a young man was discovered near present day Árpás village (Árpás-Dombiföld, Szérűskert, Hungary) in Győr-Moson-Sopron County. The deceased was laied on its back, in a shallow grave with North-South orientation, among the ruins of the Roman Mursella municipium. Grave goods included golden belt buckle, golden trouser and boot-buckles, sabretache with iron knife and forceps. Further findings were jar, glass cup, large bronze bowl, cattle leg bone, sheep sacrum and a golden coating of a probably wooden animal sculpture. The jar and glass cup have Roman parallels, but the funerary custom, costume accessories and animal sculpture have hun age eastern steppe analogies. The finding was dated to the second half of the 5th century AD[1], when the Huns ruled Pannonia.

**Singeorgiu de Mures-Kerekdomb-Grave 1 (Cx. no.41)**

During the construction of the bypass route between Corunca-Ernei (Mureş County, Romania), in the course of preventive excavations, the Archaeological Department of the Mureş County Museum discovered a Roman village and a necropolis dating from the Migration Period (according to the funerary offerings and the funeral rites), near Sângeorgiu de Mureş. The excavations brought to light 19 houses, 15 pits and 3 single-graves. According to the archaeological material the settlement can be dated to the second half of the 3rd century and the beginning of the 4th century, and the funerary offerings date the cemetery to the end of 4th century and the beginning of 5th century.

The graves were discovered on the upper part of the second terrace of the Terebici River, situated in line. Two graves were located inside the village. Grave no.1 was a partial horse burial oriented North-South, typical for the Migration Period (Huns, Alans, Goths etc.), located inside the settlement, trenched in a Roman period house [2]. Unfortunately, the grave was robbed and the right side of the human skeleton and a significant part of the horse skeleton was destroyed. The preservation of human remains was relatively good, it belonged to a 25–29 years old male with artificially deformed Mongoloid (Eastern-type) skull. The funerary offerings included a grey jug, a bone comb with cap, decorated with concentric circles and bronze plate, four silver bolts, iron knife, golden dagger gripe decorated with almandine and Cornish stone insets. The other burial (grave no. 3 Cx. no 58) was completely robbed, just the mold of the human skeleton in secondary position and remains of a wood construction on the grave`s grounds could be observed.

**Kecskemét-Mindszenti-dűlő-RL 11/2785.**
During preventive excavation of the expansion of the Mercedes factory on the outskirts of Kecskemét town (Bács-Kiskun County, Hungary) in 2017, near a late Sarmatian-Hunnic age settlement a lonely nomadic niche grave was found, containing a wealthy young male with artificially strongly deformed Mongoloid skull[3]. Grave orientation was North-South, grave goods included a dagger-sword, crescent shaped gold hair ring, a decorated gold leaf, which was probably the decoration of the knife's handle, a belt buckle, another buckle which was suspended the sword, two boot buckles and a Murga-type pottery. All of the buckles were made of silver and plated with gold, which show Hunnic analogies. The grave can be dated back to the Hunnic period on the basis of its findings. The rite of this burial and the findings differ from the traditions of the Sarmatians, who lived in this area during this period. The richness of the grave goods refer to the high social rank of the deceased. The presence of this man suggests that after the arrival of the Huns into the Carpathian Basin, they appointed one of their nobilities as leader of this late Sarmatian community. The deceased could belong to this nobility.

***Avar age burial sites:***

**Dunavecse-Kovacsos dűlő:** During preventive excavation of the M8 motorway on the outskirts of Dunavecse town (Bács-Kiskun County, Hungary) two early Avar period graves (feature 701 and 702) were found by Andrea Lantos 50 m from each other in 2004–2005. The feature 701 was a NW–SE orientation prestigious male grave, who was buried with belt decorated with pressed silver mounts, sword, bow and quiver [4]. A Byzantine jug was put near the head of the dead. The horse harness (stirrups, bit and head and breast collar and breeching) was put on the chest of the dead, the saddle decorated with bone mount was imposed on his foot. The grave was dated to 630–650/660 AD. (CSB).

**Fajsz-Garadomb**: Eight early Avar-age graves arranged in two irregular and loose line was excavated on the outskirts of Fajsz village (Bács-Kiskun County, Hungary) by Mihály Kőhegyi in 1962. The cemetery, mostly composed of armed men's graves, was the burial site of a large family or genus belonging to the middle class or leadership of Avar society. The skeletons were taxonomically Mongoloid. The deads were buried at 630–650/660 AD.
In the grave 4 lied a Mongoloid mature male skeleton; the man wore silver earrings and a belt decorated with pressed silver pseudo buckle mounts [5]. His bow and quiver with arrowheads was put into the grave. The dead was buried at circa 630 AD. (CSB)

**Kiskőrős-Vágóhídi dűlő**: 67 graves arranged in seven burial-groups of a larger Avar-age cemetery were excavated on the outskirts of Kiskőrös town (Bács-Kiskun County, Hungary) between 1934–1938 [6]. The population of the cemetery taxonomically was Mongoloid. Based on the numerous prestige objects and the gold decorate plates of the clothing and shrouds it seems to be probable, that the cemetery is the richest Middle Avar-age cemetery of the Carpathian Basin, which was used by the rich community of the power elite. The cemetery was dated between 660–720 AD.
In the grave no. I a Mongoloid mature (Sayanic) man was buried with a head-gear decorated with gold mounts and a belt decorated with round shape mounts in the middle with glass fittings. His sword was kidnapped from the grave. A silver snapper calyx was put near the dead. The dead was buried between 660–700 AD. (CSB)

**Kunbábony**: The richest known Avar-age grave was found in the border part called Bábony of Kunszentmiklós town (Bács-Kiskun County, Hungary) by Elvira H. Tóth and Attila Horváth, in 1971 [7]. Two armed man were buried near each other.
In the grave 1 was buried a man who died at the age of 60-70. His skull was Mongoloid type (Baikal type). The old man wore gold earrings, belts decorated with gold mounts. His swords, knifes, quiver and bow decorated with gold plates was put into the grave. Between the folded bones were found a gold jug, vessels decorated with gold mounts and a big amphora. dead was covered with shroud decorated gold plates. The Avar leader was buried on a bed decorated with gilded plates, which was covered with a coffin cap. 211 pieces of gold finds were found in the robbed male grave, the total weight of which was 2.33 kg. The grave-goods were dated to 630–660 AD., the burial was around 660 AD. (CSB)

**Kunpeszér-Felsőpeszéri út, Homokbánya**: 32 graves were excavated on the outskirts of Kunpeszér village (Bács-Kiskun County, Hungary) by Elvira H. Tóth [8], in which 15 graves were dated to the early Avar period, the others were buried in the 8$^{th}$ century AD. The early Avar graves were located on the large area, this was not a cemetery, but a loose burial site. Most of the dead were taxonomically Mongoloid [9]. The male graves were rich, they were buried with belt decorated silver and gilded mounts, swords decorated with gold and silver plates, bow, quiver and arrowheads. The female graves were less rich. In the graves were found simple burial attachments.
In the graves 6 and 30/B were buried adult mans. In the grave 6 were found belts decorated with silver and gilded bronze rosette shape mounts, bow and quiver made by birch bark and ten arrowheads. The man buried in the grave 30/B, wore belt decorated with silver round shape mounts; bow, quiver and sword decorated with P shaped silver ears were put into his grave. The graves were buried between 630–660 AD. (CSB)

**Makó-Mikócsa-halom**: During preventive excavation of a factory on the outskirts of the Makó town (Csongrád County, Hungary) a complete early Avar-age cemetery contained 251 graves was excavated by Csilla Balogh, in 2009–2010 [10]. On the basis of the burial customs (NE–SW orientation, catacomb and niche grave, large number of the partial sacrificial animals), who joined to the Avars in the Eastern European steppe region the cemetery seems to be used by a population of Eastern European origin [11]. Most of the population was Europid, some skulls had Mongoloid feature. The cemetery was used by the community between 568-630/650 AD.
In the niche grave 56/58 was buried a Mongoloid young man with belt decorated mounts, bow and arrowheads. Into the pit was put a horse with horse harness. The grave was dug between the last third of the 6$^{th}$ century and the first third of the 7$^{th}$ century.
In the niche grave 218/227 was buried a mature man from Europid taxon origin with belt and shoes decorated with metal mounts. In his grave were found a sword decorated with silver plates, a bow, a quiver with arrowheads and some lamella of an armor. Beside the right leg of the dead, tools for wood, bone/horn and metal processing led (saw, rasp etc.) and in the crucible made by iron plate were found some semi-finished bone bow-application. Based on the unique find-collection in Eurasia and other bone features, the man buried in the grave seem to have been also a master of. The grave can be date to the end of the 6$^{th}$ century of the beginning of the 7$^{th}$ century. (CSB)

**Petőfiszállás**: During preventive excavation of the construction of the M5 motorway on the outskirts of the Petőfiszállás village (Bács-Kiskun County, Hungary) a lonely rich armed male grave was discovered by Csilla Balogh and Erika Wicker [12]. In the grave a Mongoloid type (Saian type) 40-45 years old man was buried. His ranking belt were decorated with pressed gold mounts, his weapon belt with pressed silver round shaped with gold inlay. He wore gold earrings. In the grave were found a sword decorated with gold plates, quiver decorated with bone mounts, arrowheads and a bow. The grave was dated between 630–650/660 AD. (BCS)

**Pitvaros-Víztározó** 225 graves of the late Avar cemetery on the outskirts of the Pitvaros village (Csongrád County, Hungary) was excavated between 1993–1996 by Lívia Bende [13]. On the basis of the special burial customs the community, which has used this cemetery, seems to have been the descendants of the population of the Eastern European origin known in the early Avar period. The small community, which has opened the cemetery, stretched from the western part of the Transtisza region to the area inside the rivers around the middle of the 7th century. The community kept their special burial customs all the way (catacomb graves, partial sacrificial animals etc.). The cemetery was used between 650/660 and the end of the 8th century.
The skull of the Europid type mature man buried into the catacomb grave 72 was slightly distorted. The burial customs, primarily the catacomb grave is a characteristic of the population of the Eastern European origin, but the man wore the Meroving-type belt decorated with metal inlay, which was supposed to be a gift from a Meroving cultures community of the Transdanubia [14]. The grave was dated to the turn of the 7th-8th centuries. (BCS)

**Szegvár-Oromdűlő**: On the outskirts of the Szegvár village (Csongrád County, Hungary) 500 graves of the large Avar-age cemetery were discovered by Gábor Lőrinczy, between 1980 and 1997 [15]. On the basis of the special burial customs (NE–SW orientation, complex grave types as niche and catacomb graves, numerous partial sacrificial animals etc.) the cemetery is connected to the population of Eastern European origin, which joined to the Avars in the Eastern European steppe region. The excavated part of the cemetery was used from the end of the 6th century to the middle of the 7th century.
In the catacomb grave 81 an adult man (his skull was Mongoloid type) led, who was buried according to the burial customs of his community, but the man wore the belt decorated with cast Alpine-type mounts [16]. This belt was most likely a gift to the wearer.
In the niche grave 540 a man led too, into the grave the partial horse with horse harness was put. (BCS–LG)

**Székkutas-Kápolnadűlő**: 534 Avar-age graves were discovered on the outskirts of the Székkutas village (Csongrád County) by Katalin B. Nagy between 1965 and 1987 [17]. The catacomb graves and the partial sacrificial animals were characteristic of the cemetery. The cemetery was used from the middle of the 7th century to the beginning of the 9th century.
The grave 51 was a catacomb grave, the grave 239 was simple pit grave, but both were so-called horse-tool funeral, that is into the grave was put just the horse harness. In the graves were buried mature mans. On the basis of the Gatér-type belt mounts, the grave 51 seems to have dug rather at the end of the 630 to 650/660 period The grave 239 was dug later, in the second half or the third quarter of the 7th century on the basis of the belt mounts killed from plate. (CSB)

***Conqueror cemeteries:***

Description of the Karos-Eperjesszög I-II-III, Kenézlő-Fazekaszug I-II and Sárrétudvari-Hízóföld cemeteries were given in [18,19].

**Magyarhomorog-Kónya-domb**
This small cemetery is located at the northern border of present day Magyarhomorog village in Hajdú-Bihar County, in the eastern part of the Hungarian Great Plain. It was established in the 10th century on a small plateau on the eastern shore of the swamps of the Sebes Körös River and belongs to the a short-lived quarters type of the early Hungarians. The cemetery was excavated by István Dienes and László Kovács [20,21]. In the 17 tombs there were 11 men, three women and three infants buried in three rows of graves. Four graves (1, 9, 16, 23) were typical partial horse burials of the early Hungarians containing horse cranium with leg bones and harness objects. 7 men and the oldest child (12-14 years old) were buried with parts of archery equipment. The possibly richest man’s grave (9) was robbed, and into an

another man's tomb (15) later a body of a wolf was dug. Decades later, at the beginning of the 11th century, a large village cemetery of 523 graves was built on the same ridge.

1. Tomka, P. Az árpási 5. századi sír (The Grave of Árpás from the 5th century). in *Arrabona Múzeumi közlemények 39 /1-2* (ed. Tóth, L.) 161–188 (Győr-Moson-Sopron megyei Múzeumok Igazgatósága, 2001).
2. Gál, S. S. A Hun Age burial with artificial cranial deformation from Sîngeorgiu de Mureş - 'Kerekdomb.' in *Proceedings of the First International Conference of the Török Aurél Anthropological Association, 2016.* (ed. Gál, S. S.) 43–53 (Mega Publishing House, Cluj‑Napoca, 2016).
3. Kovacsóczy, B., Mozgai, V., Bajnóczy, B., Szabó, M. & Tóth, M. Anyagvizsgálat és készítés-technológiai megfigyelések a Kecskemét-Mindszenti-dűlőn előkerült hun kori sír nemesfémleletein. in *Hadak Útján, A népvándorláskor fiatal kutatóinak XXVIII konferenciája* 10 (Hansági Múzeum, 2018).
4. Lantos, A. Kora avar kori lószerszámos temetkezés Dunavecsén –Early Avar burial with a set of harness at Dunavecse. in *in nostra lingua Hringe nominant – Tanulmányok Szentpéteri József 60. szültésnapja tiszteletére* (eds. Csilla, B., Zsolt, P., Balázs, S. & Zsuzsanna, Z.) 29–38 (MTA Bölcsészettudományi Kutatóközpont, Kecskeméti Katona József Múzeum, 2015).
5. Balogh, C. & Kőhegyi, M. Fajsz környéki avar kori temetők II. Kora avar kori sírok Fajsz-Garadombon. — Awarenzeitliche Gräberfelder in der Umgebung von Fajsz. Frühawarenzeitliche Gräber von Fajsz-Garadomb. in *Móra Ferenc Múzeum Évkönyve – Studia Archaeologica 7* (eds. Bende, L., Lőrinczy, G. & Szalontai, C.) 333–363 (2001).
6. László, G. Études archéologiques sur'historie de la société des Avars. in *Archaeologica Hungarica 34* (1955).
7. Tóth, E. H. & Horváth, A. *Kunbábony: das Grab eines Awarenkhagans*. (Museumsdirektion der Selbstverwaltung des Komitats Bács-Kiskun, 1992).
8. H.Tóth, E. Korai avar vezetőréteg családi temetője a kunbábonyi kagán szállásterületén. in *Múzeumi kutatások Bács-Kiskun megyében* (ed. Sztrinkó, I.) 10–20 (1984).
9. Marcsik, A. A kunpeszéri avar kori széria humán csontanyagának feldolgozása (Felsőpeszéri út-Homokbánya). – Human bone material of the Avarian Age series from Kunpeszér (Felsőpeszéri út-Homokbánya). in *„In terra quondam Avarorum…" Ünnepi tanulmányok H. Tóth Elvira 80. születésnapjára. Archaeologica Cumanica 2* (eds. Somogyvári, Á. & V. Székely, G.) 175–190 (2009).
10. Balogh, C. Régészeti adatok az avar kori íjak készítéséhez – A Makó, Mikócsa-halomi kora avar kori temető 61. szerszámmellékletes sírja — Archäologische Angaben zur Herstellung der awarenzeitlichen Reflexbögen – Das Grab 61 des frühawarenzeitlichen Gräberfeldes Makó. in *Régészeti tanulmányok Nagy Margit tiszteletére – Relationes rerum – Archäologische Studien zu Ehren von Margit Nagy* (eds. Korom, A., Balogh, C., Major, B. & Türk, A.) 583–599 (2018).
11. Balogh, C. Orta Tisa Bölgesi'nde Doğu Avrupa Bozkir Kökenli Göçebe Bir Topluluğa Ait Mezarlik (Makó, Mikócsa-halom, Macaristan) – A Cemetery Belonging to an Nomad Community of Eastern Europe Steppe Origin in the Middle Tisza Region (Makó, Mikócsa-halom, Hungary). *Art Sanat* **7,** 53–70 (2017).
12. Balogh, C. & Wicker, E. Avar nemzetségfő sírja Petőfiszállás határából – Das awarenzeitlichen Sippenhäuptingsgrab von Petőfiszállás. in *Thesaurus Avarorum. Ünnepi kötet Garam Éva 70. születésnapjára. — Archaeological Studies in Honour of Éva Garam* (eds. Rácz, Z. & Vida, T.) 551–580 (2012).
13. Bende, L. Temetkezési szokások a Körös–Tisza–Maros közén az avar kor második felében. in *Studia ad Archaeologiam Pazmaniensia 8* 66–121 (2017).
14. Bende, L. Tausírozott díszű övgarnitúra a pitvarosi temetőből. – Tauschierte Gürtelgarnitur im awarischen Gräberfeld von Pitvaros. in *A Móra Ferenc Múzeum Évkönyve – Studia Archaeologica 6* 199–217 (2000).

15. Lőrinczy, G. Frühawarenzeitliche Bestattungssitten im Gebiet der Grossen Ungarischen Tiefebene östlich der Theiss. Archäologische Angaben und Bemerkungen zur Geschichte der Region im 6. und 7. Jahrhundert. in *Acta Archaeologica Academiae Scientiarum Hungaricae 68* 137–170 (2017).
16. Alpi típusú övgarnitúra a szegvár-oromdűlői 81. sírból. – Alpine-type belt set from Szegvár-Oromdűlő, grave 81. in *Zalai Múzeum 14* 137–167 (2005).
17. B.Nagy, K. A székkutas-kápolnadűlői avar kori temető. in *A Móra Ferenc Múzeum Évkönyve – Monographia Archaeologica 1* (eds. Bende, L. & Lőrinczy, G.) (3003).
18. Neparáczki, E. *et al.* Genetic structure of the early Hungarian conquerors inferred from mtDNA haplotypes and Y-chromosome haplogroups in a small cemetery. *Molecular Genetics and Genomics* 1–14 (2016). doi:10.1007/s00438-016-1267-z
19. Neparáczki, E. *et al.* Mitogenomic data indicate admixture components of Central-Inner Asian and Srubnaya origin in the conquering Hungarians. *PLoS One* (2018). doi:10.1371/journal.pone.0205920
20. Kovács, L. A Kárpát-medence honfoglalás és kora Árpád-kori szállási és falusi temetői. Kitekintéssel az előzményekre. in *A honfoglaláskor kutatásának legújabb eredményei. Monográfiák a Szegedi Tudományegyetem Régészeti Tanszékéről* (eds. Wolf, M. & Révész, L.) 511–604 (Szegedi Tudományegyetem Régészeti Tanszék, 2013).
21. Kovács, L. Előzetesen a magyarhomorog-kónya-dombi 10–12., századi temetőről. in *Népek és kultúrák a Kárpát-medencében. Tanulmányok Mesterházy Károly tiszteletére* (eds. Kovács, L. & Révész, L.) 481–501 (2016).

**Summary of anthropological data: NA = lack of data**

| LabID | Sample site/grave or accession number | Age | Age | Anthropological type | Y- Haplogroup |
|---|---|---|---|---|---|
| **Hun period** | | | | | |
| Hun/1 | Singeorgiu de Mures/1 | V. century | 25–29 | Mongolid, artificially deformed skull | Q1a2 |
| Hun/2 | Kecskemét-Mindszenti-dűlő/2785 | V. century | 18-20 | artificially strongly deformed skull with Mongolid features | R1b1a1b1a1a1 |
| Hun/3 | Árpás-Szérűskert/1 | V. century | 25-30 | Mongoloid (broad-faced) with Europid characters | R1a1a1b2a2 |
| **early Avar period** | | | | | |
| FGD/4 | Fajsz-Garadomb/4 | 630–650/660 | NA | Mongoloid | C2 |
| PSZ/1 | Petőfiszállás/1 | 630–650/660 | 40-45 | Mongoloid, Sayanic type (broad-faced, brachykran) | G2a |
| SzO/540 | Szegvár-Oromdűlő/540 | 600–650/660 | 18-20 | NA (fragmented) | I1 |
| SzO/81 | Szegvár-Oromdűlő/81 | 600–650/660 | 30-35 | Europid (Cromagnoid-B-x) | N1a1a1a1a3 |
| KFP/6 | Kunpeszér-Felsőpeszér/6 | 630–650/660 | 26-35 | NA (fragmented) | N1a1a1a1a3 |
| KFP/31 | Kunpeszér-Felsőpeszér/30B | 630–650/660 | 23-39 | NA (fragmented) | N1a1a1a1a3 |
| SzK/51 | Székkutas-Kápolnadülő/51 | 630/650–660 | 40-59 | fragmented skull with Mongolid features | N1a1a1a1a3 |
| KB/300 | Kunbábony/300 (**khagan**) | 630–650/660 | 60-70 | Mongoloid, Baicalic type with some Europid features | N1a1a |
| MM/58 | Makó-Mikócsa/56-58 | 568–630 | 18-39 | Mongoloid | N1a1a |
| MM/227 | Makó-Mikócsa/218-227 | 568–630 | 40-59 | Europid | R1a1a1b2a |
| DK/701 | Dunavecse-Kovacsos dűlő/701 | 630–650/660 | NA | NA | R1a1a1b2a |

| | | | | | |
|---|---|---|---|---|---|
| **middle and late Avar period** | | | | | |
| PV/72 | Pitvaros/72 | 650/660–700/710 | 40-59 | NA (fragmented), slightly artificially deformed skull | C2 |
| KV/3369 | Kiskőrös-Vágóhídi dűlő/3369 | 650/660–700 | 40-59 | Mongoloid (Sayanic) | N1a1a1a1a3 |
| SzK/239 | Székkutas-Kápolnadülő/239 | 650/660–700/710 | 40-59 | fragmented skull with Mongolid features | E1b1b1a1b1a |
| **Conqueror period** | | | | | |
| K1/13 | Karos I/13 | 895- mid Xth c. | 40-59 | Europid (Cromagnoid-A) | E1b1b |
| K1/1438 | Karos I/1438 | 895- mid Xth c. | 18-39 | Europid (Eu-t-p) | J1 |
| K1/1 | Karos I/1 | 895- mid Xth c. | 18-39 | NA | N1a1a1a1a4 |
| K1/10 | Karos I/10 | 895- mid Xth c. | 40-59 | NA | R1a1a1b1a2b |
| K1/3286 | Karos I/3286 | 895- mid Xth c. | 40-59 | Europid (Cromagnoid-A) | R1a1a1b2a2 |
| K2/6 | Karos II/6 | 895- mid Xth c. | 60- | Europid (Eu-t-p) | E1b1b1a1b1a |
| K2/33 | Karos II/33 | 895- mid Xth c. | 15-17 | NA | G2a2b |
| K2/26 | Karosc II/26 | 895- mid Xth c. | 18-39 | NA | I1 |
| K2/16 | Karos II/16 | 895- mid Xth c. | 60- | Europid (Cromagnoid-B) | I2a1a2b |
| K2/52 | Karos II/52 (**leader**) | 895- mid Xth c. | 40-59 | Europid (Armenoid-t-p) | I2a1a2b |
| K2/51 | Karos II/51 | 895- mid Xth c. | 40-59 | NA | N1a1a1a1a4 |
| K2/29 | Karos II/29 | 895- mid Xth c. | 18-39 | NA | N1a1a1a1a2 |
| K2/61 | Karos II/61 | 895- mid Xth c. | 60- | Europo-Mongoloid (p-t) | R1a1a1b2a2 |
| K2/36 | Karos II/36 | 895- mid Xth c. | 40 – 59 | Europid (CrC-p) | R1a1a1b1 |
| K2/41 | Karos II/41 | 895- mid Xth c. | 40 – 59 | Europo-Mongoloid (p-CrC) | R1a1a1b1a2b |
| K2/18 | Karos II/18 | 895- mid Xth c. | 60- | NA | R1a1a1b1a2b |
| K3/12 | Karos III/12 | 895- mid Xth c. | 40 – 59 | Europid (CrC-p) | I2a1a2b |
| K3/1 | Karos III/1 | 895- mid Xth c. | 40 – 59 | Europid (p-t) | R1b1a1b1a1a1 |
| K3/13 | Karos III/13 | 895- mid Xth c. | 40 – 59 | NA | R1b1a1b1a1a1 |
| K3/3 | Karos III/3 | 895- mid Xth c. | 60- | NA | R1b1a1b1a1a1 |
| KeF1/10936 | Kenézlő-Fazekaszug I/10936 | 895- mid Xth c. | 18-39 | Europo-Mongoloid | Q1a |
| KeF2/1025 | Kenézlő-Fazekaszug II/1025 | 895- mid Xth c. | 18-39 | East-Mediterranean | R1b1a1b |
| KeF2/1027 | Kenézlő-Fazekaszug II/1027 | 895- mid Xth c. | 40 – 59 | NA | N1a1a1a1a2 |
| KeF2/1045 | Kenézlő-Fazekaszug II/1045 | 895- mid Xth c. | 18-39 | Europo-Mongoloid | N1a1a1a1a2 |
| MH/15 | Magyarhomorog/15 | Xth c. | 40-45 | Europid (Cromagnoid-B) with some Mongoloid characters | I2a1a2b |
| MH/16 | Magyarhomorog/16 | Xth c. | 40-45 | Europo-Mongolid Turanid (Cromagnoid-B) | I2a1a2b |
| MH/9 | Magyarhomorog/9 | Xth c. | 40-45 | Europid (Cromagnoid-B-x) | I2a1a2 |
| SH/41 | Sárrétudvari–Hízóföld/41 | Xth c. second half | 18-39 | Europid | R1b1a1b1a1a2b |
| SH/81 | Sárrétudvari–Hízóföld/81 | Xth c. second half | 18-39 | Europid | J2a1a |
